# Network diffusion for scalable embedding of massive single-cell ATAC-seq data

**DOI:** 10.1101/2021.03.05.434093

**Authors:** Kangning Dong, Shihua Zhang

## Abstract

With the rapid development of single-cell ATAC-seq technology, it has become possible to profile the chromatin accessibility of massive individual cells. However, it remains challenging to characterize their regulatory heterogeneity due to the high-dimensional, sparse and near-binary nature of data. Most existing data representation methods were designed based on correlation, which may be ill-defined for sparse data. Moreover, these methods do not well address the issue of excessive zeros. Thus, a simple, fast and scalable approach is needed to analyze single-cell ATAC-seq data with massive cells, address the “missingness” and accurately categorize cell types. To this end, we developed a network diffusion method for scalable embedding of massive single-cell ATAC-seq data (named as scAND). Specifically, we considered the near-binary single-cell ATAC-seq data as a bipartite network that reflects the accessible relationship between cells and accessible regions, and further adopted a simple and scalable network diffusion method to embed it. scAND can take information from similar cells to alleviate the sparsity and improve cell type identification. Extensive tests and comparison with existing methods using synthetic and real data as benchmarks demonstrated its distinct superiorities in terms of clustering accuracy, robustness, scalability and data integration.

**Availability:** The Python-based scAND tool is freely available at http://page.amss.ac.cn/shihua.zhang/software.html.

## Introduction

Cell type-specific genomic regulation is driven by the binding of transcription factors (TFs) in accessible genomic regions. Thus, chromatin accessibility can be used to identify *cis*-regulatory elements and directly depict cellular identity ^[1]^. Single-cell Assay for Transposase-Accessible Chromatin using sequencing (Single-cell ATAC-seq or scATAC-seq) has enabled genome-wide profiling of chromatin accessibility at single-cell resolution and can thus reveal epigenetic heterogeneity at cellular level ^[2, 3]^.

However, clustering of single cells based on scATAC-seq data is a challenging task due to their massive dimensionality and extremely sparse nature (**Supplementary Figure S1**). For example, a recent study produced an unprecedented large-scale scATAC-seq dataset containing ~140k cells in a single study ^[4]^. One can expect that massive scATAC-seq data will be available in the near future. On the other hand, the feature dimension of scATAC-seq data is about 10 times larger than that of single-cell RNA-seq data. Thus, clustering methods for scATAC-seq data must be computationally efficient to satisfy the basic requirement of scalability and high-dimensionality. Moreover, sparsity is intrinsic to single-cell epigenomic data due to the low DNA copy number; that is, only 0, 1 or 2 reads can be captured at any genomic locus within a diploid genome. Normally, because of the limited number of captured reads, only 1–10% of expected accessible regions (i.e. peaks) could be detected per cell ^[5]^. Hence, computational analysis of scATAC-seq data is also greatly affected by the curse of “missingness”.

Existing dimension reduction and feature representation methods developed for scATAC-seq data can be roughly divided into three categories. The first category is based on known epigenomic features. For example, chromVAR ^[6]^ aggregates mapped reads into ‘cistromes’ (based on *k*-mer or motif enrichment) and measures their gain or loss of chromatin accessibility. The second category adopts natural language processing methods to treat cells as documents and accessible regions as words. Latent semantic indexing (LSI) ^[7]^ utilizes the frequency–inverse document frequency to process the binarized accessibility matrix and then reduces its dimensionality using principal component analysis (PCA). cisTopic employs the latent Dirichlet allocation topic model to generate low-dimensional topic-cell distributions from the binary epigenomic data ^[8]^. However, these two methods do not consider the “missingness” in scATAC-seq data, and cisTopic uses collapsed Gibbs sampling for distribution inference, which makes it computationally intensive. SnapATAC ^[9]^ and scasat ^[10]^ convert the accessibility matrix into a cell-by-cell similarity matrix using the Jaccard distance and extract informative features from it. However, the dense similarity matrix may be difficult to store and ill-defined due to high-dimensionality and extreme sparsity of scATAC-seq data ^[11]^. The third category is deep learning-based methods. For example, SCALE employs a deep generative model variational autoencoder to learn the low-dimensional latent representation ^[11]^. This generative model recovers missing values and is scalable to relatively large-scale datasets. But the hyper-parameters of deep learning-based methods are very hard to determine. Besides, a recently developed method named APEC converts sparse scATAC-seq data into a dense matrix by combining features with similar patterns into groups named as accesson ^[12]^. Therefore, a simple (i.e. few parameters), fast and scalable approach is urgently needed to analyze scATAC-seq data with massive cells, address the “missingness” and accurately categorize cell types.

To this end, we developed a simple and scalable dimension reduction method scAND (**<u>sc</u>**ATAC-seq data **<u>A</u>**nalysis via **<u>N</u>**etwork **<u>D</u>**iffusion). scAND starts by binarizing the peak-by-cell accessibility data matrix, which meets its near-binary nature, and directly constructs an accessibility bipartite network from the binarized matrix that an edge indicates whether the peak is accessible in the cell (**Figure 1A**). Then scAND adopts a simple and fast network diffusion method to alleviate the impact of sparsity and learn meaningful representations that preserve both local and global structure of the network. We evaluated scAND in terms of visualization and clustering on 18 synthetic datasets and 11 real datasets. Compared with existing approaches, scAND showed superior performance on the synthetic data, especially at low coverage ratio or high noise level. As for the 11 real datasets, they can be divided into two classes, depending on whether protein labeling was performed as an aid in identifying cell states. scAND substantially outperformed other methods on both types of datasets (**Figure 1B**; **Supplementary Figure S2, S3**). Three case studies (**Figure 2-4**) revealed that scAND accurately recovered the expected cell types, depicted the potential developmental trajectory and was robust to data sparsity. In addition, we also observed that the diffusion process can take information from similar cells to perform imputation. Importantly, another important merit of scAND was that it can also extract the lowdimensional representation of peaks. We employed it to integrate scATAC-seq datasets from different conditions or different platforms effectively. Last but not the least, two additional large-scale datasets and two massive simulated datasets demonstrated that scAND is fast and scalable.

**Figure 1.**
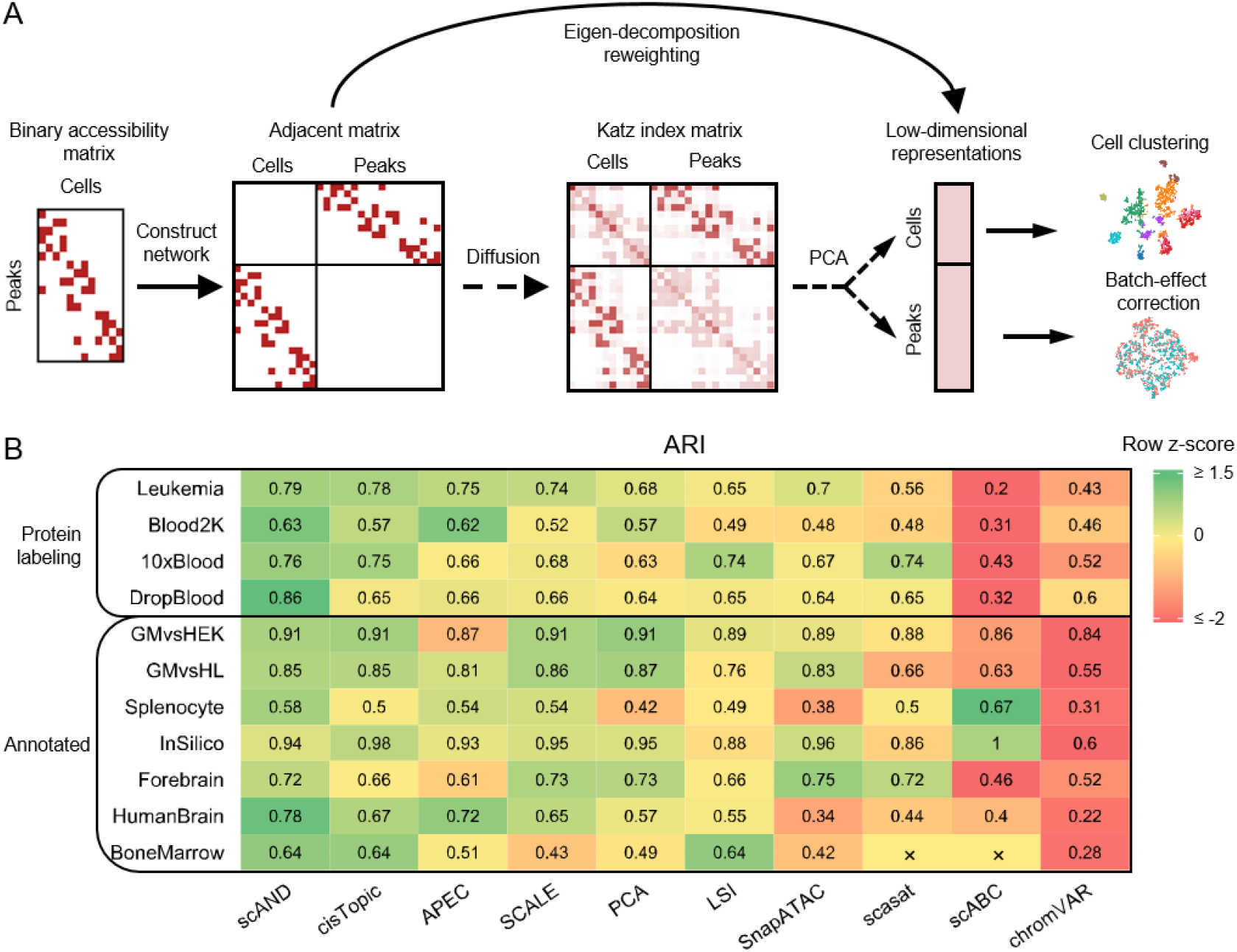
Overview of scAND (A) and its superiority to nine existing methods (B). (A) scAND first binarizes the scATAC-seq data and constructs the cell-peck bipartite network. Then scAND employs the eigen-decomposition reweighting strategy to obtain the PCA embedding of the Katz index matrix without calculating it directly for clustering and visualization. (B) Clustering performance evaluated by Louvain clustering in terms of ARI. scAND was compared with nine existing methods (columns) on 11 publicly available scATAC-seq datasets (rows). The datasets were divided into two group based on whether protein labeling was performed to aid annotation. The colors represent z-scores calculated from ARIs in the corresponding dataset. All methods were sorted by the mean ARI score in all datasets. scasat and scABC both failed on the BoneMarrow dataset due to the large number of cells.

**Figure 2.**
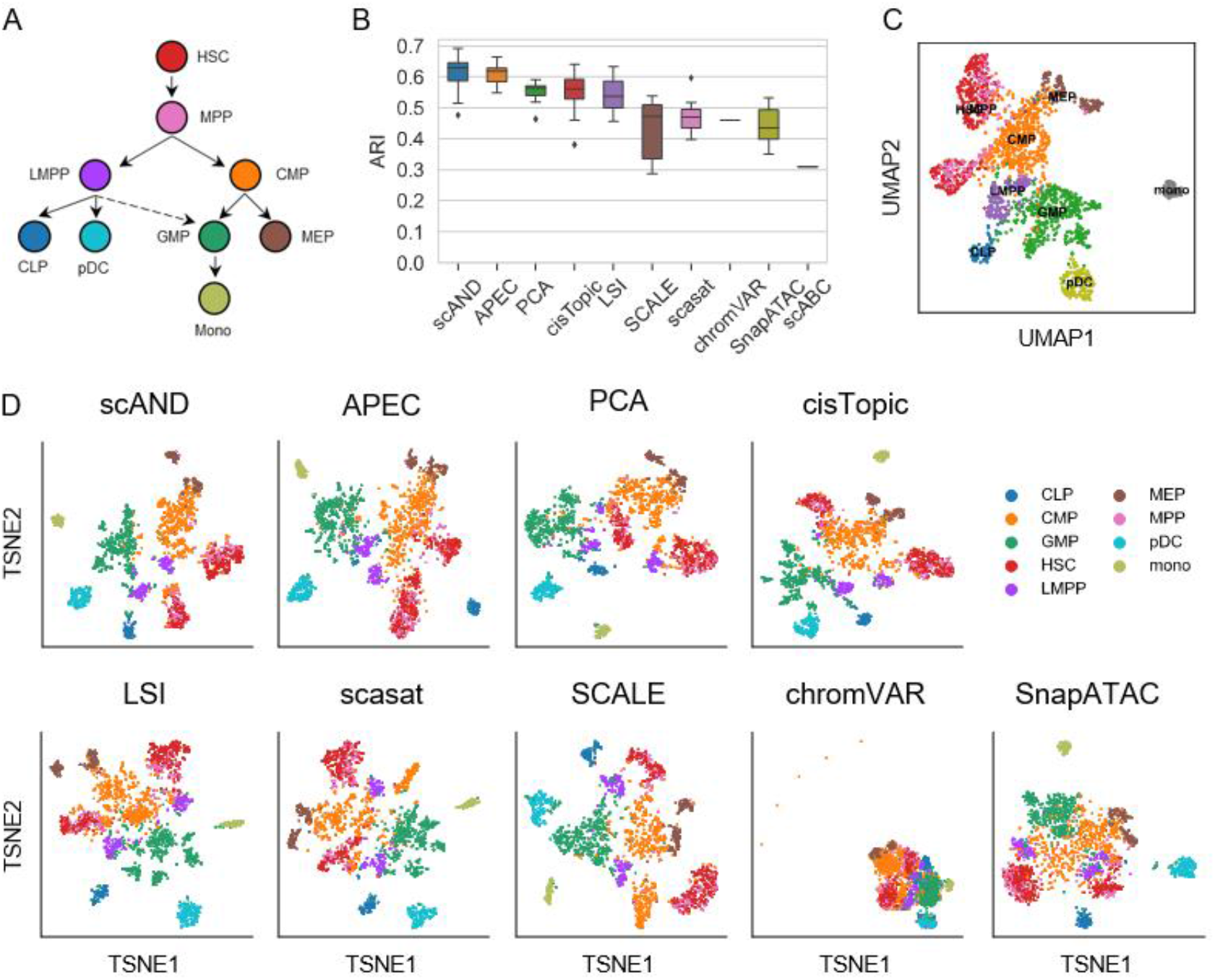
Comparison of the clustering performance and developmental trajectory of scAND in the Blood2K dataset. (A) Developmental roadmap of cell types in the Blood2K dataset. (B) Boxplots of ARIs to quantitatively measure the clustering performance of different methods with different parameters. (C) Developmental trajectory on the UMAP visualization map revealed by scAND. (D) t-SNE visualization map of the extracted features of different approaches. scABC obtained the clustering directly without using Louvain clustering, and thus there is no t-SNE visualization map for it.

## Materials and Methods

### Data

We collected 13 publicly available scATAC-seq datasets as benchmarks to demonstrate the effectiveness of scAND. 11 of them were used to evaluate the clustering accuracy, and two additional large-scale datasets were used to examine the scalability (**Supplementary Figure S1**).

- Leukemia ^[13]^: this dataset contains a mixture of HL-60 lymphoblastoid cells (HL60), monocytes and lymphoid-primed multipotent progenitors (LMPP) from healthy donors, purified leukemia stem cells (LSC) and blast cells from two patients with acute myeloid leukemia (AML).
- Blood2K ^[14]^, 10xBlood ^[15]^, DropBlood ^[4]^: these three datasets are mixtures of FACS-sorted populations from the hematopoietic system and perform protein labeling to identify subpopulations. In particular, the Blood2K dataset contains nine different cell types from the hematopoietic lineage. The 10xBlood dataset was profiled using the Chromium platform (10x Genomics) and contains CD56^+^ natural killer (NK) cells, CD14^+^ monocytes, CD19^+^ B cells, two CD8^+^ T cell subtypes and four CD4^+^ T cell subtypes. The DropBlood dataset was profiled using a droplet microfluidic device and contains CD56^+^ NK cells, CD14^+^ monocytes, CD19^+^ B cells, CD4^+^ T cells, CD8^+^ T cells and CD34^+^ cells (hematopoietic stem and progenitor cells).
- InSilico ^[3, 16]^: this dataset was merged from six individual scATAC-seq data separately profiled on different cell lines.
- GMvsHEK ^[3]^, GMvsHL ^[3]^, Splenocyte ^[17]^, HumanBrain ^[18]^, Forebrain ^[19]^, BoneMarrow ^[4]^: labels of these datasets were annotated through computational approaches. The GMvsHEK and GMvsHL datasets respectively contain cells from two commonly-used cell lines. The Splenocyte dataset was derived from mouse splenocyte cells (after red blood cells removal). The HumanBrain dataset was derived from human adult brain cells, and the Forebrain dataset was obtained from P56 mouse forebrain cells. The BoneMarrow dataset was derived from human bone marrow cells.
- Atlas ^[7]^, BM_13W ^[4]^: these two large-scale datasets were used to examine the effectiveness and scalability of methods. The Atlas dataset constructs a mouse single-cell atlas of chromatin accessibility, which includes ~80,000 cells from 13 adult mouse tissues. The BM_13W dataset contains 136,453 bone marrow cells, of which 60,495 cells are normal and the remaining 75,958 cells are in response to stimulation.

### Preprocessing

For the InSilico data, following the preprocessing of scABC ^[16]^ and SCALE ^[11]^, we only kept peaks which were accessible in ≥10 cells with ≥2 reads. For datasets with ≥200,000 peaks (Blood2K and 10xBlood), we filtered the count matrix to only keep peaks which are accessible in at least 1% cells. Note that we kept all the peaks of the HumanBrain dataset because it was extremely sparse (only a few hundred peaks left after filtering). As for the remaining datasets, we kept all the peaks.

### scAND

scAND consists of three major steps (**Figure 1A**). First, scAND directly constructs an accessibility network to represent the scATAC-seq data. Second, scAND performs network diffusion using the Katz index ^[20]^ to overcome its extreme sparsity. Network diffusion helps to smooth the sparse data, amplify the associations between cells and capture both the local and global structure of a network ^[21, 22]^. Third, scAND employs PCA to learn low-dimensional representations from the Katz index matrix. The computational complexity of calculating the dense Katz index matrix is unacceptable. We adopted an efficient eigen-decomposition reweighting strategy ^[23]^ to obtain the PCA results without calculating the Katz index matrix directly. Specifically, these steps were detailed as follows:

#### Network construction and normalization

scAND directly constructs an accessibility network from the binarized scATAC-seq data, where a node represents a peak or a cell, and an edge represents whether a peak is accessible in a cell. In order to deal with the intrinsic sparsity of single-cell epigenomic data, the weights of edges are all set to 1. Let **A**_0_ be the adjacency matrix of the network and scAND applies a symmetric square root graph normalization to alleviate the coverage bias in single-cell experiments. The normalized adjacency matrix **A** is computed as follows:

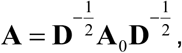

where **D** is a diagonal matrix whose diagonal element *D_ii_* represents the sum of the *i*th row of **A**_0_.

#### Network embedding via network diffusion

Katz index ^[20]^ is one of the most widely-used diffusion-based similarity measure. It is defined as follows:

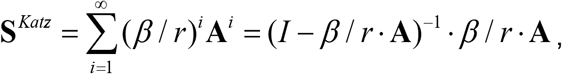

where **A** is the adjacency matrix of the network, *r* is the spectral radius of **A** and *β* ∈ (0,1) is a decay parameter (*β* <1 to ensure convergence ^[24]^). Actually, 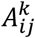 is equivalent to the sum of weights of walks of length *k* from node *i* to node *j.* Thus, the Katz index matrix propagates and smooths the network by taking information from all the nodes that have more than one path between them. In addition to alleviating the sparsity of scATAC-seq data, the cell-by-cell submatrix (peak-by-peak submatrix) of the Katz index matrix encodes the global similarity information among cells (peaks).

However, it is not feasible to directly calculate the Katz index matrix for high-dimensional scATAC-seq data. In order to reduce the computational complexity, scAND adopts the eigen-decomposition reweighting strategy to achieve the network embedding ^[23]^ to directly obtain singular value decomposition (SVD) of the Katz index matrix, which is based on the following matrix theory.

#### Theorem

(Eigen-Decomposition Reweighting). If *x* is an eigenvector of **S**, and *λ* is its associated eigenvalue. For any polynomial function *F*(*x*), *x* is also an eigenvector of *F*(**S**), and *F*(*λ*) is its associated eigenvalue.

Because *S^Katz^* is a symmetric matrix, its SVD decomposition and eigen-decomposition are equivalent. To calculate the top-*d* SVD results of *S^Katz^*, it first calculates the top-*l* (*l* ≥ *d* where *l* satisfies that at least *d* eigenvalues are positive) eigen-decomposition [**Λ, X**] of adjacency matrix **A**, and calculates the reweighted eigenvalues **Λ**^*Katz*^ = *F*(**Λ**), where *F*(*x*) = (*I* – *β*/*r* · *x*)^-1^ · *β*/*r* · *x*. Then it sorts **Λ**^*Katz*^ in descending order of the absolute value and selects the top-*d* of them. These top-*d* reweighted eigenvalues and their associated eigenvectors are the top-*d* eigen-decomposition of Katz index matrix ***S**^Katz^*. So, we can directly obtain the top-*d* SVD results of ***S**^Katz^* without calculating ***S**^Katz^*. Let’s denote the top-d SVD results of ***S**^Katz^* as [**U,∑,V**]. The PCA embedding can be obtained by multiplying the square-root of **Σ** into **U**:

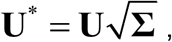

The U* contains both the low-dimensional representations of cells and peaks. The cell representations capture the characteristics of cells, which can be visualized in the low-dimensional space using t-distributed stochastic neighbor embedding (t-SNE) and used to detect cell populations with various clustering methods like Louvain or K-means. The peak representations are in the same latent low-dimensional space with those of cells, and can be further used to correct batch-effects.

Moreover, it is worth noting that the first principal component may mainly relate to the sequencing depth rather than biologically variability in some cases ^[7, 25]^. Some approaches such as LSI ^[7]^ and scasat ^[10]^ removed the first principal component when performing downstream analyses. However, we believe that directly removing the first principal component may lose some information. Thus, we recommend to perform *L*_2_ normalization on cell representations, i.e. dividing each representation by its *L*_2_ norm. We found this normalization indeed helped to improve clustering performance (**Supplementary Table S1**).

#### Parameter settings of scAND

scAND has two hyperparameters: the diffusion parameter *β* and the dimension *d* of PCA embedding. The diffusion parameter *β* relates to the sparsity issue and the emergence of new cell subpopulations with the changing of it. We introduced a leave-out imputation strategy for its selection. Explicitly, we randomly set 10% entries to 0 and calculated the L2 distance between the true data and the imputed one with scAND. We tried *β*s from 0 to 1 in steps of 0.05 for all the datasets, and chose the value with the smallest distance (**Supplementary Figure S4**).

As for the dimension of PCA embedding *d*, we adopted the “elbow” plot to select the number of principle components *d* from 10, 20, 30, 40 or 50 with similar principle in Seurat ^[26]^. We also evaluated the clustering performance of [*d*-10, *d*+10] in steps of 1 and the results is not sensitive compared to other methods (**Supplementary Figure S5**).

### Computational complexity

Computational complexity of scAND is mainly determined by the number of edges in the network (i.e. the number of non-zero fragments in data). Explicitly, for a network with *N* nodes (the sum of the numbers of cells and peaks) and *M* edges (twice the number of non-zero fragments of scATAC-seq data), computational complexity of square root graph normalization is *O*(2*M*) by using sparse matrix multiplication. Computational complexity of calculating top-*/* eigen-decomposition of the adjacency matrix is *O*(*T*(*Nl*^2^ + *Ml*)) by using iterative approaches (a Python function *scipy.sparse.linalg.eigs)* ^[27]^, where *T* is the number of iterations. Computational complexity of eigen-decomposition reweighting is *O*(*l* +*Nd*). So, the total computational complexity is *O*(*T*(*Nl*^2^ + *Ml*)). Since *l* is usually small (*l* is slightly higher than the dimension *d* and *d* << *N*), computational complexity of scAND is linearly determined by the number of non-zero fragments in scATAC-seq data. Thus, scAND is fast and scalable for the sparse scATAC-seq data.

### Clustering and visualization

We used the Louvain community detection-based clustering algorithm ^[28, 29]^ to perform cell clustering (implemented by the Python package *Scanpy* ^[30]^). The size of local neighborhood of Louvain clustering was set as 15 and the random state was set as 2019 in all cases. Unlike other clustering approaches such as K-means, Louvain clustering requires a tunable parameter “resolution” instead of the number of clusters. We used the same process as ^[5]^ to apply the binary search algorithm on “resolution” to obtain the same number of clusters as the number of labels of a dataset.

We used t-SNE ^[31]^ and UMAP ^[32]^ implemented by the Python *Scanpy* package ^[30]^ to project the low-dimensional representations to a two-dimensional space with random state as 2019. The number of neighbors for t-SNE and UMAP were set as 15 and 30 respectively.

### Evaluation of clustering performance

#### Adjusted Rand Index (ARI)

ARI measures the similarity between true labels and predicted clusters by considering both matched and unmatched assignment pairs. It is defined as follows:

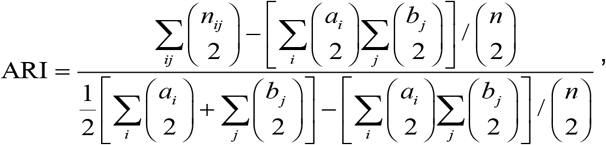

where *n_ij_* is the number of cells (that belongs to the *i*th cell type) is assigned to the *j*th cluster, *a_i_* and *b_j_* are the number of cells labeled as the *i*th cell type and *j*th cluster respectively. ARI is between 0 (random labeling) and 1 (perfectly matching).

#### Normalized Mutual Information (NMI)

NMI is used to measure the mutual dependence between two random variables. It is computed as follows:

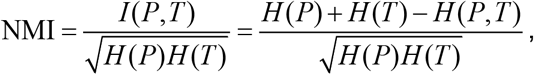

where *P* and *T* are empirical categorical distributions for predicted clusters and real cell types, *I*(*P, T*) is the mutual entropy, *H(P)* and *H(T)* are the Shannon entropies of *P* and *T* respectively, and *H(P, T)* is the joint entropy of *P* and *T.* NMI is bounded between 0 (mutual independent) and 1 (perfect correlation).

#### Homogeneity Score (HS)

A perfectly homogeneous clustering meets an important requirement: a cluster should only contain samples belonging to a single class. HS is a measure to describe the closeness of clustering results to the perfect homogeneity clustering. It is computed according to the following formula:

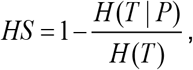

where *H*(*T*|*P*) is the conditional entropy, *H(T)* is the Shannon entropy. HS is bounded between *0* and *1,* while *1* stands for the perfectly homogeneous clustering.

### Comparison with existing methods

We compared the clustering performance of scAND with eight state-of-art approaches: PCA, cisTopic ^[8]^, APEC^[12]^, LSI ^[7]^, SnapATAC ^[9]^, scasat ^[10]^, scABC ^[16]^, SCALE ^[11]^ and chromVAR ^[6]^ (see **Supplementary Table S2** for their parameter settings).

Generally, the number of dimensions for the PCA-based methods (PCA, LSI, scasat and SnapATAC) are determined by the “elbow” point for the variance plot of the 50 largest principal components. Explicitly, the first principal component was removed for LSI and scasat in all cases. For PCA, we performed the *L*_2_ normalization process by dividing each representation with its *L*_2_ norm, which significantly improves the clustering results (**Supplementary Table S1**). For APEC, the results of different number of accessons (from 500 to 1500 in steps of 50) were shown. The number of topics for cisTopic was selected from 5 to 50 in steps of 1 using the parameter selection process of the R package *cisTopic.* As for SnapATAC, the process as recommended started by a bin-by-cell binary matrix generated through segmenting the genome into bins of uniform size. However, in order to fairly compare with other methods, SnapATAC was applied to the peak-by-cell binary matrix. Notably, to speed up for large-scale datasets, SnapATAC calculates the partial Jaccard index matrix where it only contains the similarity between all pairs of *N* cells with randomly selected *K* cells (*K* << *N*). We adopted this approximate process with *K*=2000 for large-scale datasets (10xBlood, DropBlood, HumanBrain and BoneMarrow). As for SCALE, we used the default network architecture (two-layer network with 1028-128 and 10 latent variables for GMM). We used the k-mer version of chromVAR in this paper, because its performance is generally better than the motif version in our experiments ^[5]^.

For the sake of fairness, the clustering performance measured by ARI of different parameters were explored (**Supplementary Figure S5**). Specifically, for PCA-based methods and cisTopic, results of different number of dimensions (from [*d*-10, *d*+10] in steps of 1 where *d* is the selected dimension) were shown. For APEC, results of different number of accessons (from 500 to 1500 in steps of 50) were shown. For SCALE, we conducted a grid search on the number of layers (i.e. 2, 3 and 4) and the dimension of latent features (i.e. 10, 20 and 30). The random states of all methods were set as 2019.

### Association of cell populations of scATAC-seq and scRNA-seq data

We connected cell populations of scATAC-seq and scRNA-seq data via gene set overlap analysis. We first calculated the gene activity score from scATAC-seq data using Cicero ^[33]^ and used *‘t-test’* implemented in the *scanpy* package ^[30]^ to identify differentially expressed genes. The FDR threshold for the test was set as 1% (Benjamini-Hochberg adjusted), and the log fold change threshold was set as 0.5. Then we calculated the *P* values of the Fisher’s exact test for the overlap of differential genes of scATAC-seq data and marker genes of scRNA-seq data, and the *-log*(*P-values*) were used to construct the association matrix. Finally, we calculated the z-score for each row and columns of the association matrix.

### Batch effect correction

scAND could correct batch effects by using the Canonical Correlation Analysis (CCA) algorithm on the peak representations (Figure 6A). Explicitly, for datasets with batch effects, we first run scAND on each dataset individually. Secondly, we used peak representations of the same peaks in different datasets to train the CCA model, which further projected the cell representations to the CCA space to obtain the corrected ones. The dimension of CCA model was set to 5 for all cases.

## Results

### scAND consistently improves clustering performance on synthetic data

To explore the performance and robustness of scAND, we first applied it onto 18 synthetic datasets with different read coverage ratios (2,500, 5,000 or 10,000 fragments per cell) and noise levels (from 0 to 50%; 10% by 1). In our simulation, we performed downsampling from nine FACS-sorted hematopoietic system related bulk ATAC-seq data ^[3]^ and perturbed them with random sampling in terms of different noise levels (**Supplementary Figure S6**). We compared the clustering accuracy of scAND with nine competing methods (**Methods**) in terms of ARI with dimension equaling 10 for all methods.

We found that all methods performed well on the high-coverage datasets. As for the low-coverage datasets, scAND and LSI were the only two methods that clearly separated all nine cell populations, but other methods failed to separate hematopoietic stem cells (HSCs) and multipotent progenitors (MPPs) due to high correlation between their bulk data (**Supplementary Figure S6B**; the Pearson correlation coefficient > 0.99). As to robustness to noise, scAND and LSI performed significantly better than other methods on the low-coverage datasets (**Supplementary Figure S7 and S8**). Although scAND cannot distinguish HSCs and MPPs when the noise level > 20%, the remaining seven populations were clearly separated.

### scAND demonstrates superior clustering performance on real scATAC-seq data

We further applied scAND onto 11 publicly available scATAC-seq datasets generated from different platforms (**Methods; Supplementary Figure S1**) and compared it with nine competing methods. Here, we adopted the Louvain clustering algorithm for all the methods (**Figure 1B and Supplementary Figure S2**). scAND achieved overall best performance on 9 datasets, and was almost as accurate as scABC on the InSilico dataset. As to the Splenocyte dataset, it was the densest one (16.6% non-zero fragments) among all, but all methods did not perform well due to the defect of Louvain clustering which may divide some clear cell populations into smaller ones (**Supplementary Figure S9**). To remove the impact of different clustering methods, we applied K-means clustering algorithm and still found that scAND achieved one of the best among all methods (**Supplementary Figure S3**). Notably, in the Leukemia, Blood2K, 10xBlood and DropBlood datasets, protein labeling was used to define cell types. scAND substantially outperformed other approaches on the DropBlood dataset, and was one of the best methods on the other three datasets, suggesting that scAND is superior to competing methods in uncovering biologically relevant subpopulations.

scAND can also depict the developmental trajectory and reflect the relative distance between different cell subpopulations in UMAP ^[32]^ visualization (**Supplementary Figure S10**). For example, the Blood2K dataset contains nine FACS-sorted differentiating cell types from the human hematopoietic lineage (**Figure 2A**). scAND correctly identified these cell types and depicted the expected developmental trajectory (**Figure 2B, C**). In particular, scAND and APEC were the only two methods that clustered the two subpopulations of the lymphoid-primed multipotent progenitors (LMPPs) together and depicted the multiple stages of the lineage during hematopoietic development (**Figure 2D and Supplementary Figure S11**). For the Forebrain dataset, scAND clearly separated the three previously annotated subpopulations of excitatory neuron cells (EX1, EX2 and EX3), which were close to each other. Meanwhile, we applied Louvain clustering algorithm with resolution 0.5 on the scAND representations and identified 6 subtypes of excitatory neuron cells and 4 subtypes of inhibitory neuron cells (**Supplementary Figure S12B**). We found that these subtypes well correspond to those identified from a mouse visual cortex scRNA-seq datasets ^[34]^ (**Methods**, **Supplementary Figure S12C, D**). On the contrary, as for the datasets consisting of cells from different cell lines, scAND made a significant distinction among different cell types. For example, as to the InSilico dataset consisting of cells from six different cell lines, scAND clearly separated four of them with significant distinction, and only cells from HL60 and TF-1 cell lines (both of which are erythroleukemic cell lines) were closely clustered. Another example is the Leukemia dataset, which contains both cells from distinct cell lines and similar cells from AML patients (**Supplementary Figure S13**). scAND, APEC and cisTopic were tied for the best clustering performance, where scAND showed four better separable cell populations from AML patients in the t-SNE visualization map. Specifically, scAND, APEC and cisTopic can perfectly divide the three distinct cell lines (LMPPs, HL60 and monocytes) with median AIR = 1. In terms of clustering the four cell populations from AML patients, scAND (median ARI = 0.627) outperformed APEC (median ARI = 0.578) and cisTopic (median ARI = 0.567).

Moreover, scAND was robust to data sparsity and performed better in extremely sparse data. For example, on the sparsest HumanBrain dataset, scAND achieved the best performance of clustering (**Supplementary Figure S9**). Notably, scasat and SnapATAC were not effective in this situation, suggesting that the similarity-based approaches may be ill-defined due to the sparsity.

### scAND can uncover cell subpopulations within biologically complex dataset

As we have mentioned that the BoneMarrow dataset were annotated with 15 cell populations based on LSI. scAND unexpectedly achieved competitive performance with LSI among all methods (**Figure 1B and Supplementary Figure S5**). More specifically, scAND identified two subpopulations in pre-B cells and two subpopulations in CD4^+^ T cells (**Figure 3A and 3B**). We employed Cicero ^[33]^ to infer the gene activity, and found that the gene activity of SATB1 and TNFRSF13B were differential between subpopulations of pre-B cells, and the activity of LEF1, LAX1 and TBX21 gene was heterogeneous between CD4^+^ subpopulations (**Figure 3C**). SATB1 has been reported to regulate development of hematopoietic progenitor cells ^[35]^, and the TNFRSF13B mutation impaired the B cell activation process ^[35–37]^. LEF1 and LAX1 have been found differentially expressed at resting T cells and activated T cells ^[38]^, and TBX21 regulated the CD4^+^ T cell Th1/Th2 cell fate decision ^[39]^. We also observed that the inferred activity of POU motifs such as POU2F1 using chromVAR ^[6]^, which has been found to orchestrate B cell fate ^[40]^, were heterogeneous between two subpopulations of pre-B cells (**Figure 3D**), which likely reflected the heterogeneous developmental states within pre-B cells. And the motifs of JunB, a regulatory of CD4^+^ T cell fate ^[41]^, showed differential accessibility between the two subpopulations of CD4^+^ T cells. However, the subpopulations of pre-B cell were not as clearly distinguishable in the t-SNE visualization map of other methods (**Supplementary Figure S14B**). In addition to revealing the new subpopulations, scAND could well characterize the subtypes annotated in the BoneMarrow dataset. For example, scAND clearly separated the two subtypes of monocytes, which showed different motif activity of marker TF SPI1 and CEBPA ^[4]^. For the human hematopoietic stem and progenitor cells (HSPCs), Lareau *et al.* ^[4]^ has aligned the Blood2K dataset onto these cells based on the UMAP representation. We annotated the HSPCs by using 10-nearest neighbor classifier on this UMAP representation and observed that scAND depicted two hematopoietic differentiation lineages within the HSPCs, from HSCs to monocytes and from HSCs to erythroid respectively (**Figure 3A and Supplementary Figure S14C**).

**Figure 3.**
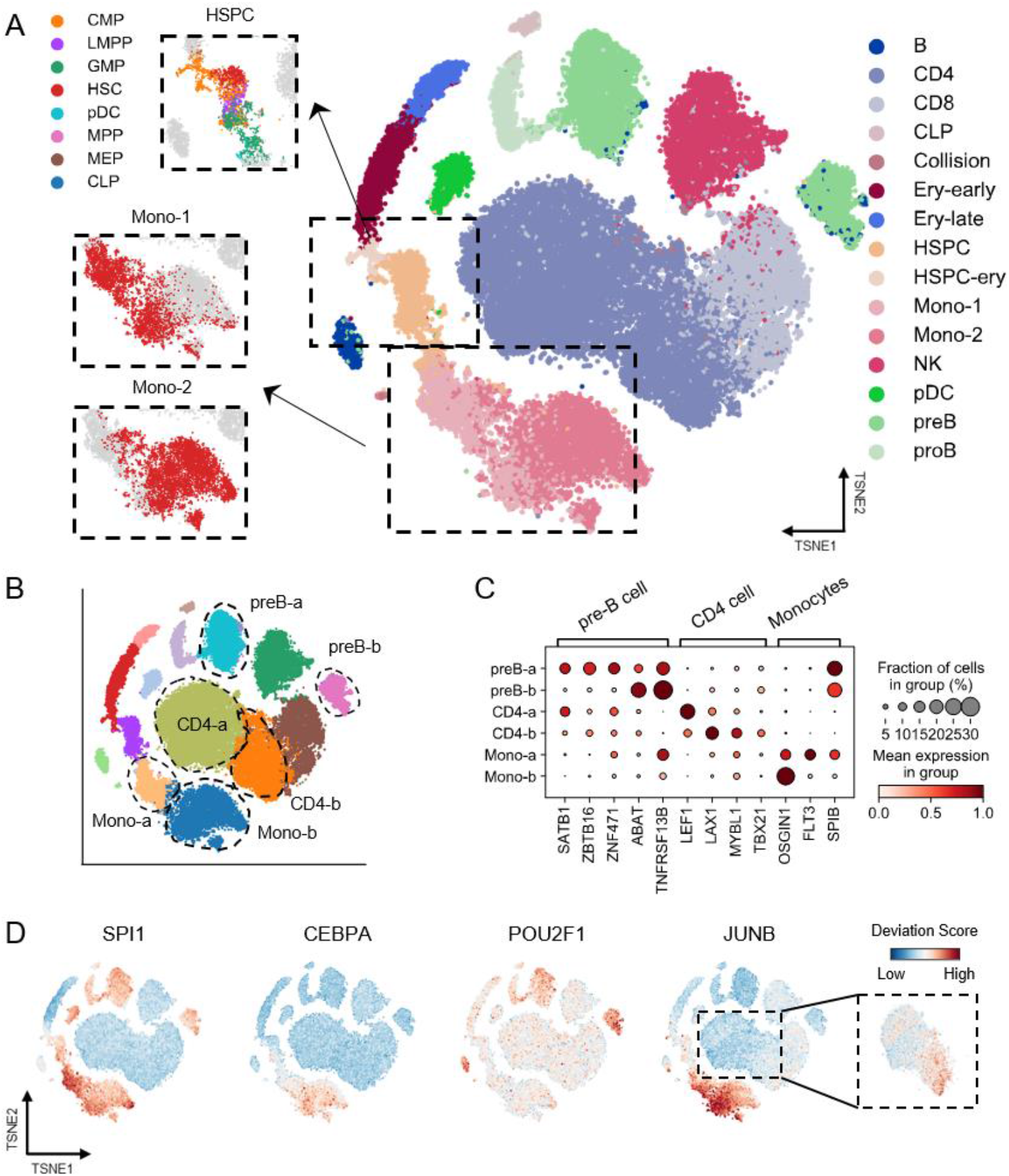
Comparison of clustering performance of scAND and other methods and their diverse regulatory heterogeneity in the human bone marrow cells. (A) t-SNE visualization of scAND representations colored by the reference cell types, including B cells, CD4^+^ T cells, CD8^+^ T cells, common lymphoid progenitors (CLP), erythroid (Ery-early and Ery-late), human hematopoietic stem and progenitor cells (HSPC), Monocytes (Mono-1 and Mono-2), natural killer cells (NK), plasmacytoid dendritic cells (pDC), pre-B cells and pro-B cells. The upper inset shows the transferred labels of HSPCs, which were obtained by the alignment with the Blood2K dataset. The lower inset shows the two monocyte clusters. (B) t-SNE visualization maps of scAND representations from the BoneMarrow dataset. The colors of dots indicate the clustering label of Louvain clustering. The dotted circles represent the identified subpopulations of pre-B cells, monocytes and CD4^+^ T cells. (C) The average Cicero score (color) and the fraction of expressing cells (circle size) of differentially expressed marker genes in pre-B cell subpopulations, CD4^+^ T cell subpopulations and monocytes subpopulations. (D) t-SNE visualization colored by TF motif accessibility scores of SPI1, CEBPA, POU2F1 and JUNB, computed using chromVAR.

scAND significantly outperforms other methods on the DropBlood dataset (median ARI = 0.859; **Figure 4A**) and is the only method that clearly separated CD4^+^ T cells and CD8^+^ T cells (**Figure 4B, C** and **Supplementary Figure S15B**). In particular, the NK cells, monocytes and B cells were divided into two subpopulations respectively, and we observed heterogeneous motif activity of cell development-related TFs across these subtypes (**Figure 4D**), such as that of TCF7L2, which is related to T cell development ^[42]^, and those of CEBPA in monocytes ^[4]^ and POU2F2 in B cells ^[43]^, indicating the different developmental states of cells (**Figure 4E**). For the three CD8^+^ T cell clusters, we annotated the resting CD8^+^ T cell cluster based on the lower motif accessibility of EOMES ^[44]^. Moreover, scAND identified 4 clusters for the CD34^+^ HSPCs, where the identification of CLP was supported by the high motif accessibility of EBF1 ^[4]^, and the other three clusters constructed a potential development trajectory from HSPC-a to HSPC-c, which is supported by the differential motif activities of hematopoietic stem cell generation marker gene GATA2 ^[45]^ and the monocyte marker gene CEBPA ^[4]^. However, the two subpopulations of B were not clearly separated in the t-SNE visualization map generated by other methods, as well as the HSPC-a and HSPC-b cells (**Supplementary Figure S15C**). And the 10xBlood dataset consists of 18,140 cells and had been annotated 9 cell types based on the protein labeling. As expected, scAND was one of the best methods in this case (**Supplementary Figure S16**). Actually, we observed that the two replications of Memory CD4^+^ T cells and Naïve CD4^+^ T cells were separated in the visualization map respectively, suggesting the potential batch effects. Although influenced by the batch effects, scAND still clearly separated cells into five major cluster consistent with the protein labeling: B cells, monocytes, NK cells, CD4^+^ T cells and CD8^+^ T cells (**Supplementary Figure S16C**). Collectively, these results showed that scAND can dissect the cellular heterogeneity and better identify subpopulations with biological interpretation.

**Figure 4.**
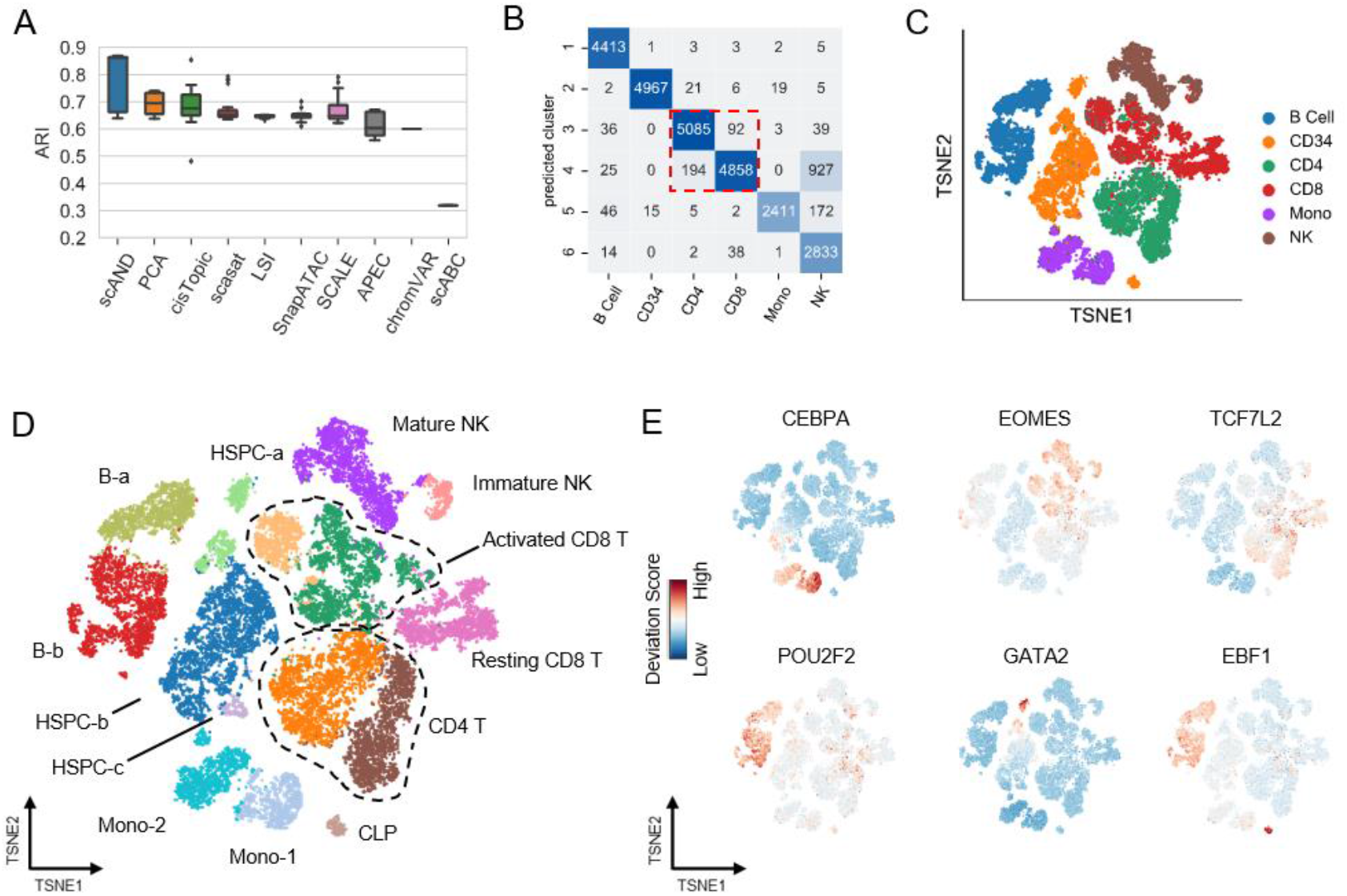
Cell subtypes detection of scAND and other methods on the DropBlood dataset. (A) Boxplots of ARIs to quantitatively measure the clustering performance of different methods with different parameters in the DropBlood dataset. (B) Confusion matrix between cluster assignments of scAND and reference cell types, which were given by protein labeling. (C) t-SNE visualization map of scAND representations colored by the reference cell types. Note that CD34 represents the CD34^+^ stem/progenitor cells (HSPCs), Mono represents monocytes, and NK represents natural killer cells. (D) t-SNE visualization map of Louvain clustering with resolution parameter as 0.5 on the scAND representations. scAND identified 15 clusters, which were annotated based on protein labeling and TF motif accessibility. (E) t-SNE visualization maps colored by TF motif accessibility scores of CEBPA, EOMES, TCF7L2, POU2F1, GATA2 and EBF1, computed using chromVAR.

### Diffusion indeed helped to reveal the regulatory heterogeneity

Next, we evaluated whether the diffusion process indeed contributed to better characterizing of the regulatory heterogeneity. We observed that the clustering performance of scAND with higher diffusion levels were generally better than those without diffusion (*β*=0) (**Supplementary Figure S17A**). More interestingly, we found that stronger diffusion could amplify the cell heterogeneity in some cases. We used two examples to illustrate it. The first example is the two pre-B subpopulations in the BoneMarrow dataset (**Figure 5A**). Compared to the result without diffusion (*β*=0), scAND divided these two subpopulations more clearly through diffusion. Another example is the HSPC-a subpopulation in the DropBlood dataset (**Supplementary Figure S17B**), which tended to be clearly separated with the increasing of diffusion level. Based on these examples, the necessity of diffusion process is evident.

**Figure 5.**
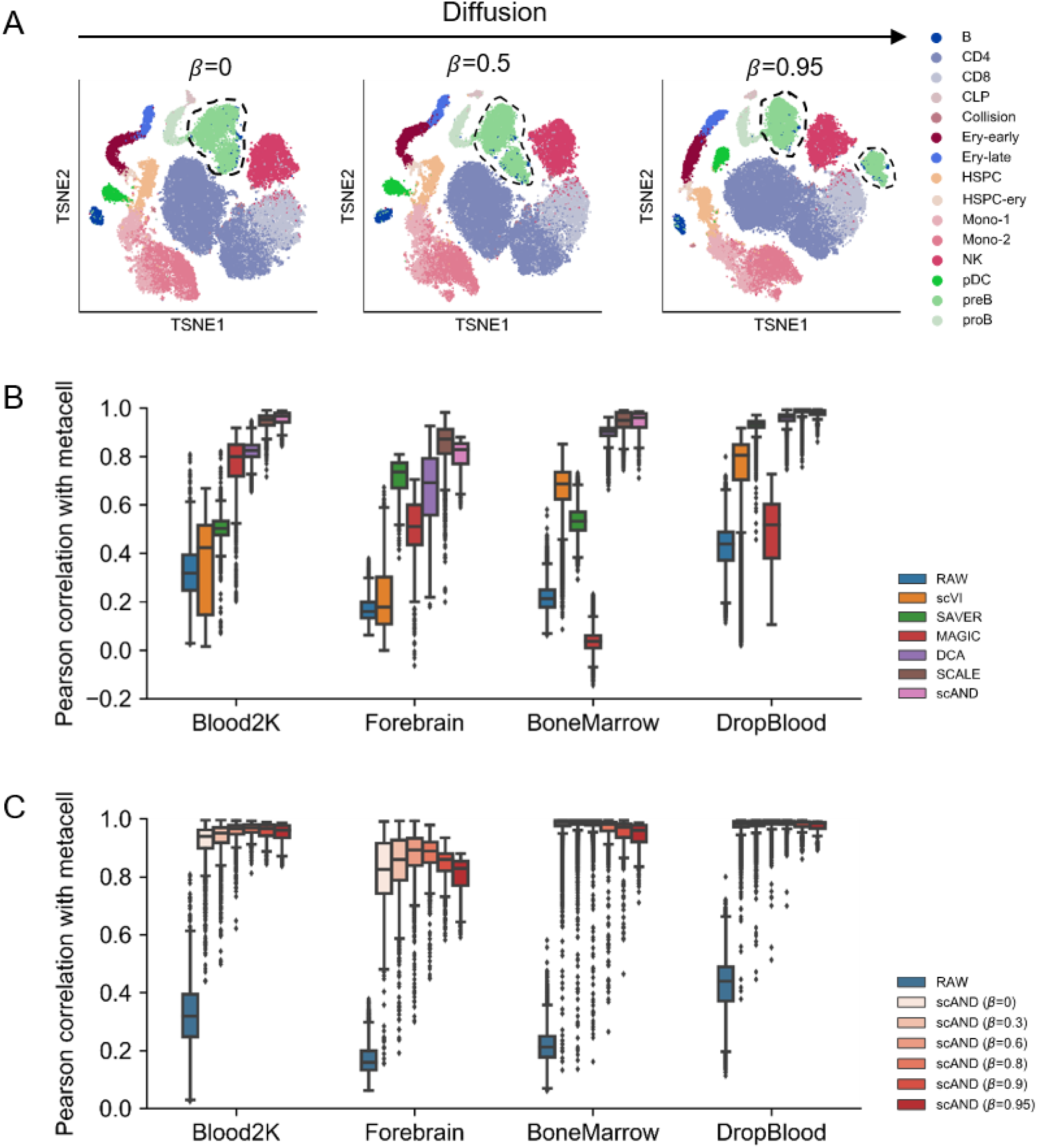
Performance evaluation of scAND with different diffusion levels. (A) t-SNE visualization maps of scAND representations generated with different diffusion parameter β on the BoneMarrow dataset. The dotted circles represent the two subpopulations of pre-B cells (see Figure 3). (B) Boxplots of cell-wise Pearson correlations between the raw data and the restored one with the meta-cell of each cell type on the four datasets. (C) Boxplots of cell-wise Pearson correlations between the raw data and the scAND restored ones using different diffusion parameter β with the meta-cell of each cell type.

An important feature of network diffusion is the ability to alleviate the sparsity by taking information from the similar cells. The diffusion process reflects the similarity relationship between cells and peaks. Thus, the restored matrix can be used as an imputed version of the raw data. Specifically, scAND first multiplies U* and V* to restore the diffused Katz index matrix and then performs the inverse operation of the normalization to get the imputed data. To approximate the ground truth accessibility of cells, we defined meta-cell by the average of all single cells from the same biological cell type ^[46]^. Generally, the correlations between raw data and resulted meta-cell are low due to the extreme sparsity ^[46, 47]^. Thus, we expected that the imputation methods could improve the correlations to aid in cell population identification. Here, we compared scAND with SCALE and four state-of-the-art scRNA-seq imputation algorithms scVI ^[48]^, SAVER ^[49]^, MAGIC ^[22]^ and DCA ^[50]^ in four datasets (**Figure 5B**). scAND and SCALE were tied for the best and achieved the highest correlations. More interestingly, as the diffusion level increasing, the correlation strengthened, and the number of cells deviated from their corresponding meta-cell decreased (**Figure 5C**), indicating the diffusion process helped to recover the potential accessibility structure of cells.

Moreover, the imputation of scAND could remove noise and enhance the cluster-specific biological signals. We performed chromVAR analysis on the imputed data with scAND and the raw data in both Blood2K and Forebrain datasets (**Supplementary Figure S18**). The t-SNE visualization map of chromVAR on the imputed data showed clearly separable clusters compared to those on the raw data, and the clustering accuracies were improved in terms of ARI accordingly (from 0.362 to 0.579 for the Blood2K dataset and from 0.411 to 0.565 for the Forebrain dataset). Many cell type-specific TFs showed much more specific patterns in the imputed results, such as the known monocyte-regulating TF ELF1 ^[51]^, hematopoietic stem cell fate FOXO4 ^[52]^, myeloid progenitors development-related TF NFIL3 ^[53]^ and lymphoid progenitor-related TF TBX21 ^[39]^.

### Peak representations of scAND can be applied to batch effect correction

scAND could obtain the representation of peaks in the same low-dimensional space of cells. We suggested a simple data alignment and batch effect correction algorithm using the peak representations (**Methods**; **Figure 6A**), and demonstrated it on a large-scale dataset BM_13W consisting of 60,495 normal bone marrow cells and 75,958 stimulated bone marrow cells ^[4]^. Cell representations of different conditions from scAND showed that cells tend to cluster under stimulation condition rather than by cell types (**Figure 6B**). After alignment and integration, the t-SNE visualization map verified that cells grouped entirely by cell types and were properly aligned across conditions (**Figure 6C, D**).

**Figure 6.**
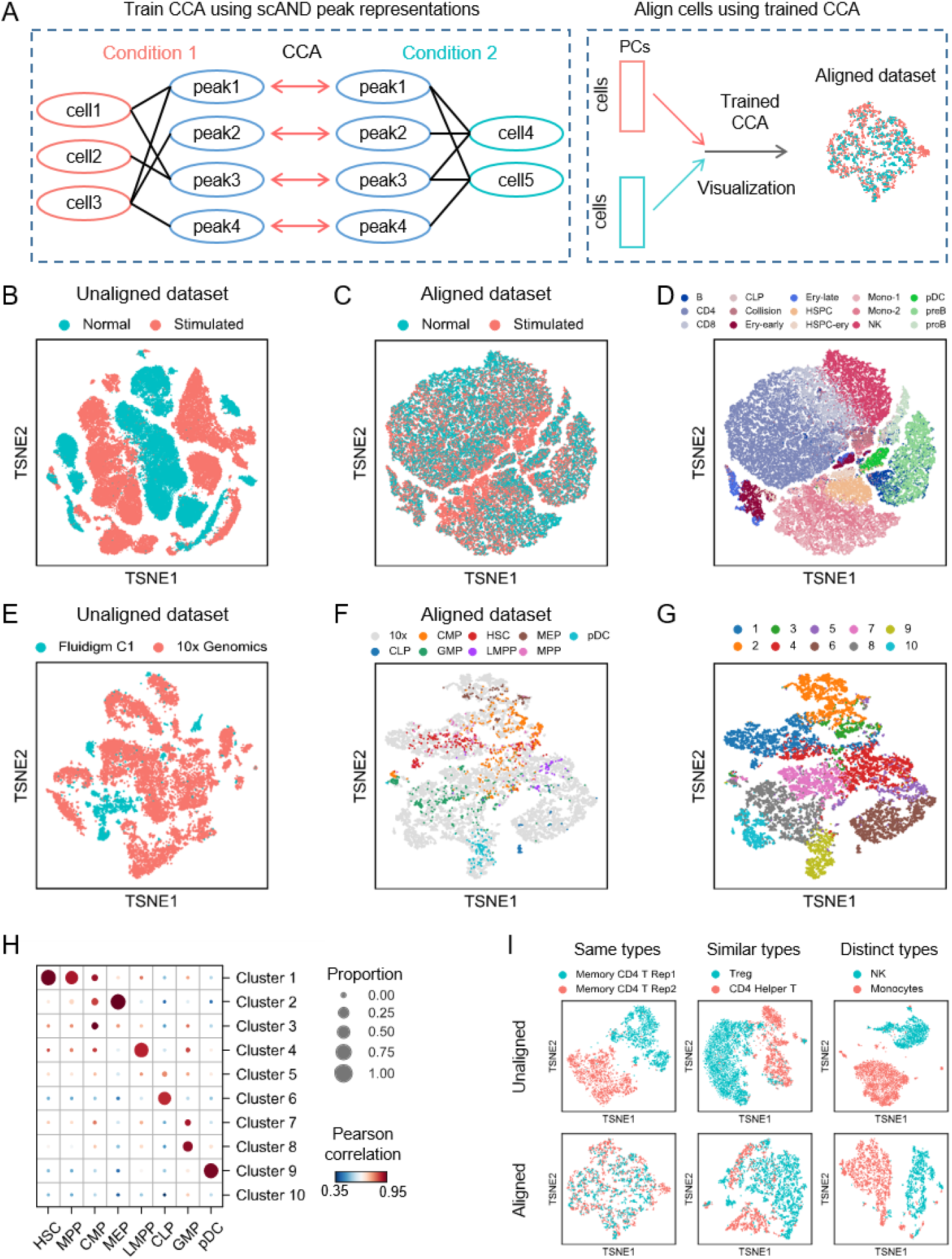
Integration of two scATAC-seq datasets using scAND peak representations. (A) Overview of the data integration strategy. (B-D) t-SNE visualization maps of human bone marrow cells split between normal (60,495 cells) and stimulated (75,958 cells), before (B) and after (C, D) integration. (E-G) t-SNE visualization maps of hematopoietic CD34^+^ cells from two different platforms, Fluidigm C1 (2,034 cells) and 10x Genomics (16,415 cells), before (E) and after (F, G) integration. The dots of (F) is colored by the cell types of scATAC-seq data from Fluidigm C1 and those of (G) is colored by the cluster results of scATAC-seq data from 10x Genomics. (H) Confusion matrix of sorted hematopoietic populations showing the proportion of each population in clusters defined in (G). The color intensity reflects the Pearson correlation between meta-cell of corresponding populations or clusters. (I) t-SNE visualization maps of three case studies indicating different batches of the same, similar, and distinct cell types respectively.

We also applied this integration method on datasets generated from different platforms: the Blood2K dataset generated from Fluidigm C1 ^[14]^ and the CD34+ populations of 10xBlood dataset generated from 10x Genomics ^[15]^. The integration strategy properly aligned cells across platforms and preserved the information of cell types in the Blood2K dataset, while directly merged combination fails to do that (**Figure 6E-G**). Importantly, the correlation of meta-cell of different populations were consistent with the proportion of each population in clusters, suggesting that this integration algorithm can well transfer information across datasets (**Figure 6H**).

An important merit of our integration strategy is that it is trained on the peak representations rather than those of cells, which takes advantages of the large number of peaks and can avoid over-correction of cells. We collected three subsets from the 10xBlood dataset with different degrees of difference: (1) two replicates of memory CD4^+^ T cell populations; (2) two similar cell types, Treg and CD4^+^ Helper T cell; (3) two distinct cell types, NK cells and monocytes. Here we compared scAND with the widely-used batch effect correction methods Harmony ^[54]^ and SeuratV3 ^[26]^, and found that the integrated results of scAND most prominently reflect the differences among cell types (**Figure 6I and Supplementary Figure S19**). Specifically, for the case contains two distinct cell types, Harmony mixed NK cells and monocytes together, Seurat integrated a small proportion of cells and scAND clearly separated these two distinct cell types. These results showed that peak representations of scAND can be applied to integrate different datasets and correct batch effects.

### scAND is scalable to massive datasets

Here we used two additional large-scale scATAC-seq datasets with more than 80,000 cells and two simulated massive datasets to demonstrate that scAND is computationally efficient and scalable. The clustering result of scAND showed a good agreement with the reference cell types inferred by the atlas study (ARI=0.803) (**Supplementary Figure S20**).

Theoretically, the computational complexity of scAND scales almost linearly with respect to the number of non-zero fragments in scATAC-seq data (**Methods**). Compared with other methods, scAND was only slower than PCA and SnapATAC in large-scale datasets (**Table 1** and **Supplement Table S3**). While SnapATAC adopts an approximate method for large-scale data that only calculates the similarity of randomly selected *K* cells (*K*=2000 by default) with all *N* cells. When *K* was set to 5000, SnapATAC cost more running time accordingly and failed to run on the simulated dataset with 50W cells (**Table 2**). Compared to cisTopic, scAND significantly reduced the running time using the same computing resource. When dealing with the largest real dataset Atlas which included ~80,000 cells and ~400,000 peaks, scAND only cost 25 minutes and was nearly 38 times faster than cisTopic (**Table 1**). More importantly, only our Python-based tool scAND can run on the simulated dataset with 100W cells (in about 5 hours). These results indicated that scAND can handle massive datasets that one will face in the near future.

**Table 1.**
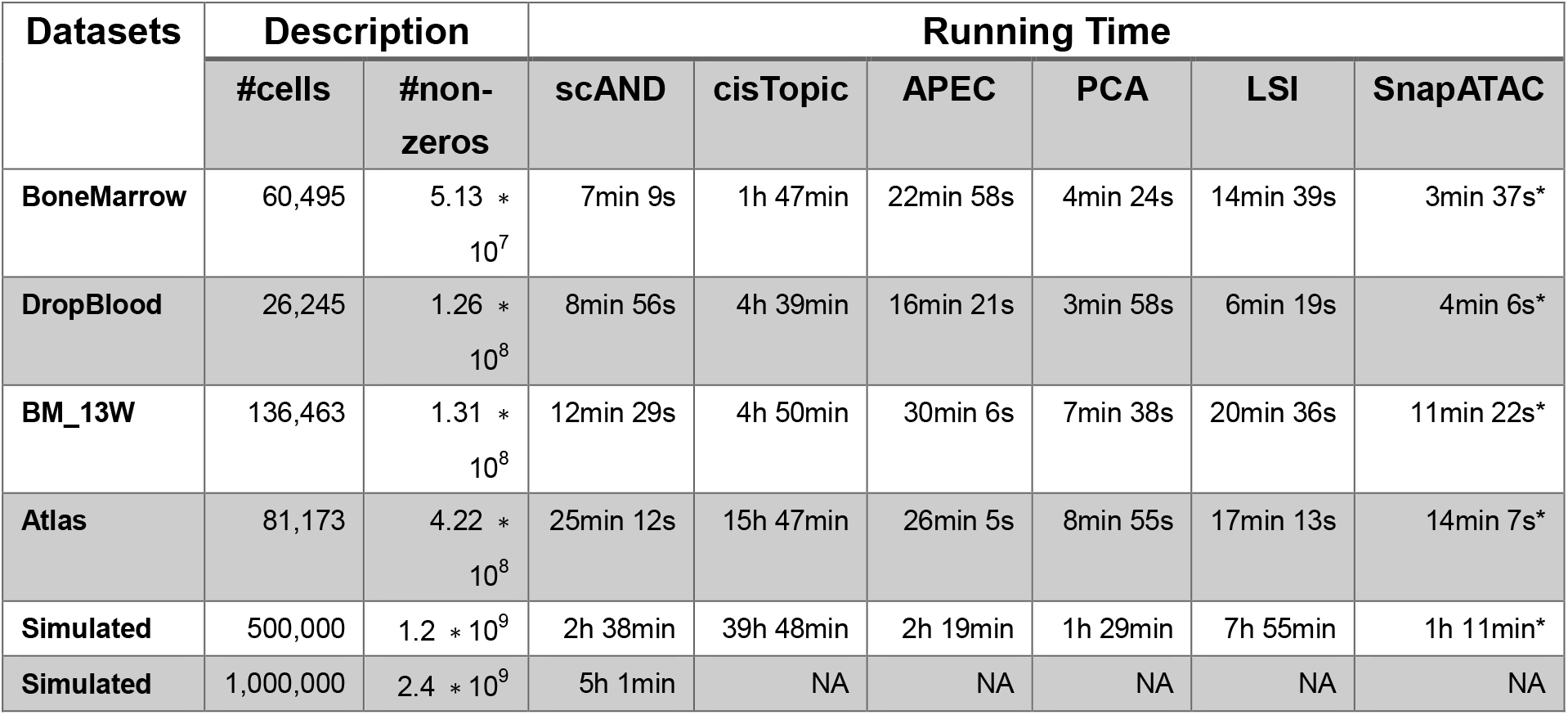
Computational time for dimension reduction using scAND and other methods on four illustrative real datasets and two massive simulated datasets. All the tests were run on a machine with 2 Intel Xeon CPU E5-2643. *indicates that SnapATAC calculates the partial Jaccard index matrix where it only contains the similarity between all *N* cells and randomly selected *K* cells (*K*=2000).

**Table 2.**
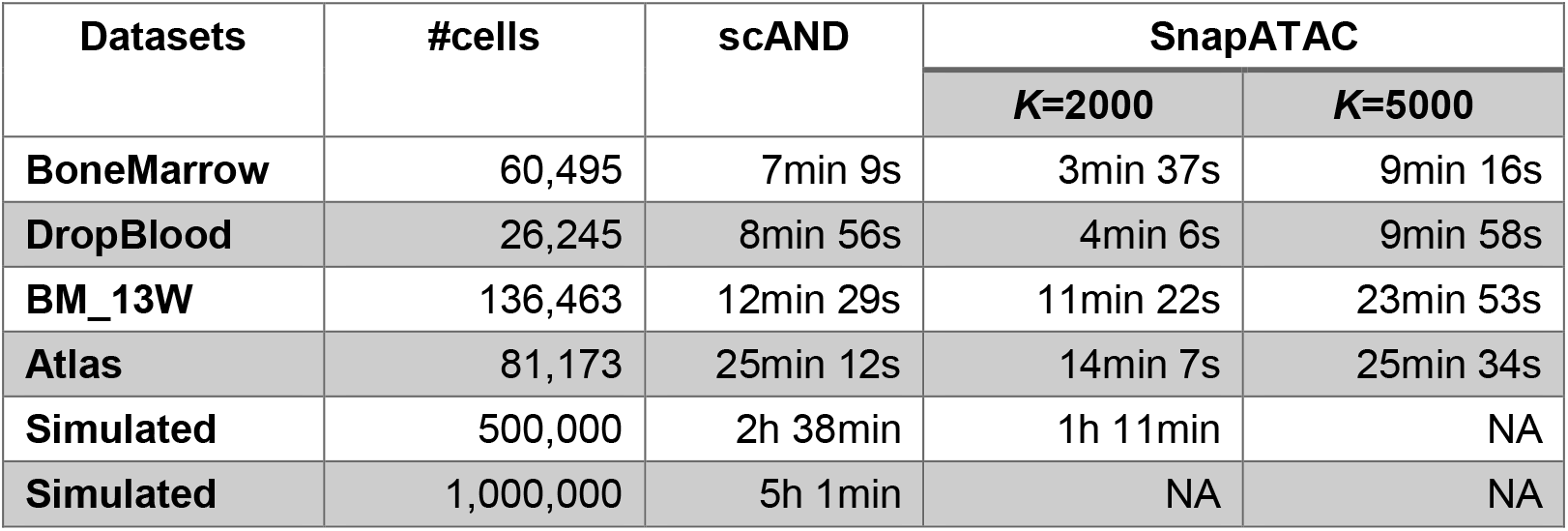
Computational time of scAND and SnapATAC on six large-scale datasets. All the tests were run on a machine with 2 Intel Xeon CPU E5-2643. SnapATAC calculates the partial Jaccard index matrix where it only contains the similarity between all *N* cells and randomly selected *K* cells (*K*=2000 or 5000).

## Discussion

The key challenges in analyzing single-cell epigenomic data is how to deal with its near-binary sparsity and massive scale. Here, we developed a simple, fast and scalable dimension reduction approach scAND for analyzing scATAC-seq data. scAND treats it as a bipartite network that indicates the accessible relationship between cells and peaks. Then the Katz index matrix is used to propagate and smooth the network, which also gathers global information of the network. scAND achieved significantly better clustering accuracy in comprehensive experiments including both 18 synthetic datasets and 11 real scATAC-seq datasets. Moreover, the visualization of scAND representations revealed the developmental trajectory and reflected the relative distance between cell types. The cell accessibility recovered by scAND showed higher correlation with their corresponding meta-cell compared to those of raw data. These results indicated that scAND representation can be applied to further downstream analysis such as developmental trajectories reconstruction and imputation to reduce the computational complexity and improve the performance.

The success of scAND is mainly attributed to the diffusion process. In the diffused Katz index matrix, missing values are restored by taking information from similar cells and peaks, and the cell-by-cell submatrix encodes the correlations between cells. These two features have been verified to aid in improve clustering accuracy in existing methods: APEC achieves state-of-the-art performance by combining the similar peaks; snapATAC learns the low-dimensional representations from cell correlation matrix. We consider that scAND utilizes the advantages of these two methods without increasing the computational complexity. Moreover, we also illustrated the ability of diffusion to impute the data, which could help to improve the performance on extremely sparse data. We observed that scAND displayed only a moderate decrease in clustering accuracy in all cases, even with the excessed data corruption rates larger than 0.6 (**Supplementary Figure S21**). In general, cisTopic was also robust with increased data sparsity; but their clustering accuracies dropped when the corruption rate > 0.6.

Consistent with the assessment paper for scATAC-seq data clustering methods ^[5]^, except scAND, APEC, LSI and cisTopic were the best approaches. However, we observed that PCA was even better than LSI and SnapATAC in terms of both clustering accuracy and time cost (**Figure 1B**), while the results of previous study ^[5]^ showed that PCA is not scalable to large-scale datasets. This difference was mainly caused by two reasons. First, there were many fast and memory-efficient PCA algorithms and implementations, including the Golub-Kahan method implemented by the Python package sklearn ^[55]^ and the IRLBA method implemented by the R package *irlba* ^[56]^. The acceleration of PCA implementations can significantly reduce the computational time when applied to sparse single-cell data with millions of cells ^[57]^. Second, performing normalization on PCA embedding is necessary. We performed *l*_2_ normalization on PCA in this work and found that this normalization process substantially improved the clustering accuracy (**Supplementary Table S1**). We believe that, since PCA-based methods have few parameters and are scalable, they are more applicable to analyzing sparse scATAC-seq data as the increasing of data size compared to deep learning-based methods.

The idea to represent a sparse data using a bipartite network in scAND is direct, and can be applied to other sparse single-cell omics data such as single-cell methylation data ^[58]^ or single-cell ChIP-seq data ^[59]^. We applied scAND onto a single-nucleus methylcytosine sequencing (snmC-seq) dataset containing neuronal populations from the human frontal cortex and mouse frontal cortex ^[60]^ (**Supplementary Figure S22**). scAND can well capture the cell subpopulations as showed in the t-SNE visualization map.

## Acknowledgments

This work has been supported by the National Key R&D Program of China [2019YFA0709501]; the National Natural Science Foundation of China [61621003]; the National Ten Thousand Talent Program for Young Top-notch Talents, the CAS Frontier Science Research Key Project for Top Young Scientist [QYZDB-SSW-SYS008] and the Shanghai Municipal Science and Technology Major Project [2017SHZDZX01].

## Author contributions

SZ conceived and supervised the project. KN designed, implemented and validated the algorithm and results. KD and SC wrote the manuscript, read and approved the final manuscript.

